# Comment on ‘Multiple Origin but Single Domestication Led to *Oryza sativa*’

**DOI:** 10.1101/299669

**Authors:** Peter Civáň, Terence A. Brown

## Abstract

In 2015, we published an analysis of rice genomic data and showed that the *japonica*, *indica* and *aus* groups of cultivated rice were independently domesticated. Our conclusions were controversial as they contradicted a previous, high-profile analysis of the same dataset, which had suggested that all of cultivated rice derives from a single origin. Although there have been attempts since 2015 to bolster the single-origin hypothesis, until recently there has been no direct rebuttal of the methodology that we used to infer multiple origins. Such a rebuttal has now been published (Choi J.Y., Purugganan M.D., 2018 Multiple origin but single domestication led to *Oryza sativa.* G3 8:797-803), but the reanalysis that is presented only supports the single origin hypothesis if phylogenetic trees that are clearly paraphyletic are interpreted as monophyletic, and furthermore addresses only one component of the evidence that we presented for multiple domestications. We caution against accepting these analyses uncritically.

---

In a recent issue of G3: Genes/Genomes/Genetics, Choi and Purugganan (2018) reanalysed the rice genomic dataset of Huang *et al*. (2012) including use of the approach for identification of putative selective sweeps devised by us (Civáň *et al*. 2015). They interpret the outcomes as supporting a single *de novo* domestication of Asian rice followed by transfer of domestication alleles to other wild populations by introgression. In contrast, we concluded that there were three separate domestications of Asian rice. The reanalysis of Choi and Purugganan (2018) is technically sound and in some respects superior to the previous two studies. Their methodology differs from ours in several ways: (i) they used more sophisticated tools for genotype reconstruction from whole genome sequencing reads, presumably yielding a data matrix of higher quality; (ii) they performed diversity scans with genomic windows of fixed physical length rather than windows with fixed number of single nucleotide polymorphisms (SNPs), (iii) they use percentile-based thresholds for identification of selective sweeps, as opposed to a fixed threshold uniform for all *Oryza sativa* groups; (iv) they used all accessions for the construction of co-located low-diversity genomic region (CLDGR) trees, rather than replacing the cultivated haplotypes with group majority consensus haplotypes. However, there are two serious flaws in the interpretation of their results which invalidate their conclusion that there was a single domestication of Asian rice.

Choi and Purugganan (2018) base their conclusion on the observation that in each of the neighbour-joining trees for the three CLDGRs carrying the *Sh4*, *Prog1* and *Laba1* genes, which are functionally related to domestication, all the domesticated rice accessions clustered together, displaying what they describe as monophyletic relationships. However it is incorrect to describe these clusters as monophyletic. For each of these genes, and for each of the examined genomic window sizes, the recovered topology is not a monophyletic *O. sativa* group but instead is paraphyletic (Figures 1B, 2A, 2B and 2C in Choi and Purugganan 2018). This is a crucial distinction, because while a monophyletic *O. sativa* clade would indeed indicate a single origin for the given genomic region in *O. sativa*, a paraphyletic group has no such implication. In each of those trees, the group containing *O. sativa* also contains many *Oryza rufipogon* genotypes: e.g. with the shortest, 40 kb, windows shown in Figure 1B there are >100 *O. rufipogon* genotypes in each of the *O. sativa* groups for the *Sh4* and *Prog1* regions, and there are ~70 such genotypes in the *Laba1* region). The paraphyly within these clusters in fact suggests that cultivated rice obtained the examined regions from multiple *O. rufipogon* individuals. This could have occurred during a single domestication, but equally could have occurred during multiple domestication processes. In agreement with the latter interpretation, we have recently documented multiple genealogical lineages of the *Sh4* and *Prog1* regions in cultivated rice (Civáň and Brown 2017).

**Figure 1.**
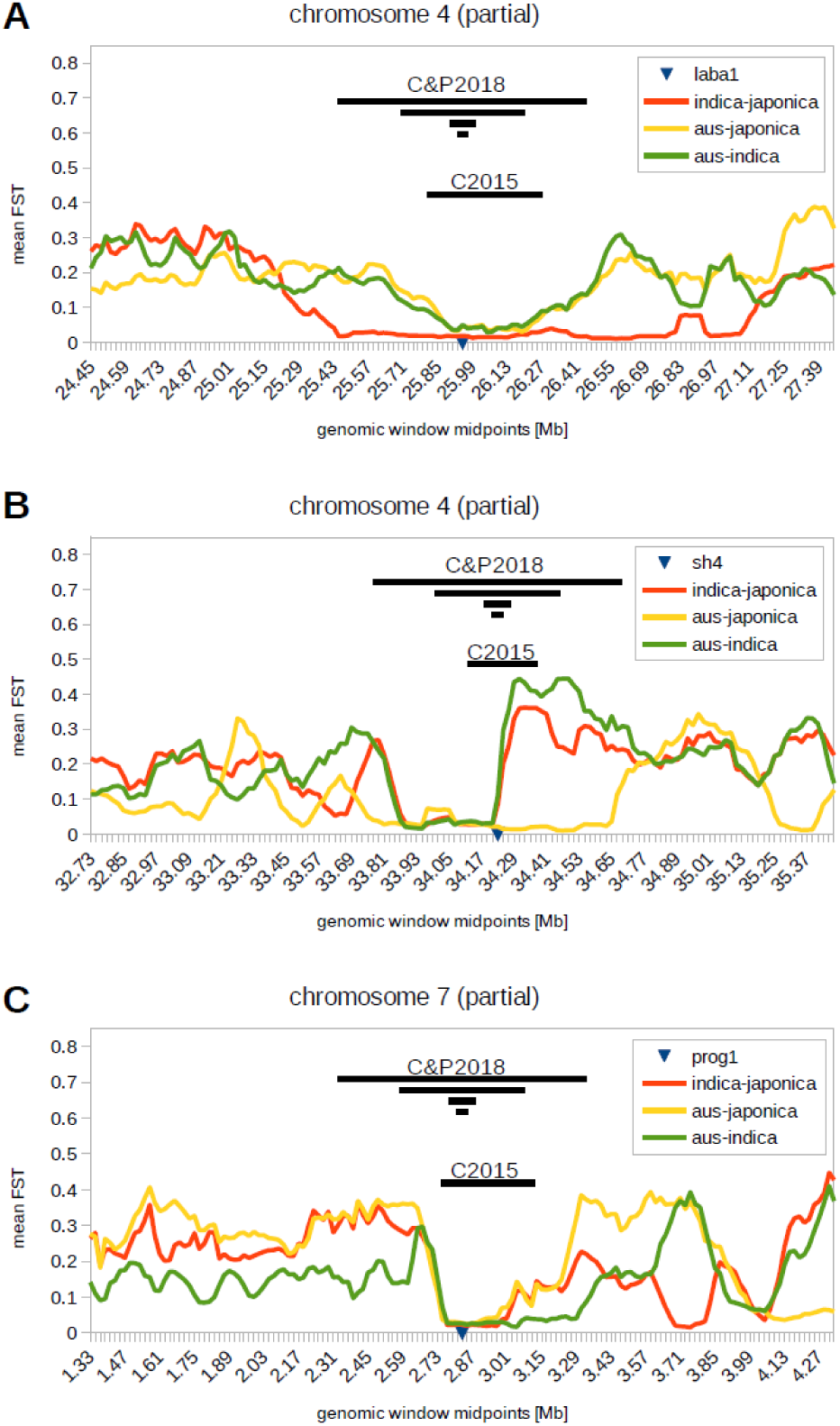
Genomic regions containing the *Laba1* (A), *Sh4* (B) and *Prog1* (C) genes. Genomic windows analysed by Civáň *et al*. (2015) and Choi and Purugganan (2018) are indicated with black bars. Graphs were produced using the 29 million biallelic SNP dataset (3,000 rice genome project, 2014) downloaded from http://oryzasnp-atcg-irri-org.s3-website-ap-southeast-1.amazonaws.com. The bed file was converted into vcf format using PLINK v1.90 (S. Purcell and C. Chang www.cog-genomics.org/plink/1.9/; Chang *et al*. 2015), selecting 283 *indica*, 154 tropical and temperate *japonica* and 124 *aus* individuals. Pairwise *F*_ST_ values were calculated with VCFtools 0.1.15 (Danecek *et al*. 2011), using 100 kb window size and 20 kb sliding steps (physical length of the IRGSP-1.0 genome assembly).

The regions surrounding the *Laba1*, *Sh4* and *Prog1* genes also appeared within the CLDGRs that we identified (CLDGR15 for *Laba1*, CLDGR16 for *Sh4*, CLDGR21 for *Prog1*, see Supplementary Figures 3l, 3m and 3r, respectively, in Civáň *et al*. 2015). If we set aside the the monophyly–paraphyly error in Choi and Purugganan (2018), their analysis of these three regions gives very similar results to those we reported. For the *Laba1* and *Prog1* regions (Figures 1A and 1C), both studies found that all cultivated rice contains similar haplotypes that cluster together. In the case of *Sh4*, we found that the consensus sequence for *indica* differs from those of *japonica* and *aus* and is more closely related to some haplotypes found in *O. rufipogon.* Choi and Purugganan (2018) argue that the *Sh4* upstream region (to the right of *sh4* in Figure 1B) is not useful for such a comparison, because the putative *Sh4*-introgression block does not exceed far upstream of the *Sh4* gene. We agree with this possibility, but excluding the *Sh4* upstream region leads again to a paraphyletic tree with similar haplotypes present in *indica*, *japonica* and wild rice. Presence of the *sh4* ‘domestication’ allele in wild rice (without phenotypic effect) has been confirmed by Sanger sequencing (Zhu *et al*. 2012) and we have shown that the causative SNP pre-dates domestication (Civáň and Brown 2017). It is therefore impossible to infer whether *indica* obtained the *sh4* allele from *japonica* or its wild progenitor, and the paraphyletic trees in Choi and Purugganan (2018) are inconclusive. These data alone cannot be used to support any domestication model.

Importantly, the scarcity of the shared sweeps (i.e. monophyletic CLDGRs) is only one part of the argument that led us to conclude that there were three independent domestications of Asian rice. Equally important is the abundance of group-specific (i.e. unshared) selective sweeps. Group-specific selective sweeps were detected in the original study (Huang *et al*. 2012) and by us (Civáň *et al*. 2015), and although Choi and Purugganan (2018) do not mention them, it is implied that they detected them too. Domestication can be viewed as a long-term selection experiment, and the signatures of group-specific selection are likely to be the signatures of separate domestications within that long-term process. Each of *indica*, *japonica* and *aus* has a unique selection profile, manifested by their different phenotypes, ecological preferences and culinary properties. Even if a few genes were transferred between the groups by introgressive hybridization, domestication exerted selection on dozens of other loci and the results we presented in Civáň *et al*. (2015) show that selection acted on different gene pools in different geographic regions and were performed by different human communities. Therefore, the conclusion of Choi and Purugganan (2018) that “*de novo* domestication appears to have occurred only once” is unwarranted.

Although Choi and Purugganan (2018) fail to interpret their trees as paraphyletic, they are aware of the observation that domestication-related haplotypes are frequently found in wild rice. Instead of interpreting this as ancestral variants that had been selected during domestication(s), they suggest a somewhat non-parsimonious explanation – that the domestication variants evolved during domestication and were subsequently transferred to wild rice by crop-to-wild gene flow. In support of this hypothesis, they mention the recent paper by Wang et al. (2017), who claim that “most modern wild rice is heavily admixed with domesticated rice”. Even though we agree that some level of gene flow is likely to have occurred in this direction, we are convinced that the scale of this problem is greatly exaggerated by Wang et al. (2017) and Choi and Purugganan (2018). As Wang et al. (2017) rightly point out, the genetic similarity of wild and domesticated rice can be due to ancestor-descendant relationship or to gene flow. They propose that a strong correlation between genetic and geographic distances in crop-wild pairs would support the latter. Subsequently, they report “a highly significant correlation (ρ=0.15, P<2.2 × 10^−16^)” and conclude this signals gene flow. This is, however, a misleading interpretation of the statistical test. In fact, although the P-value indicates that the result is extremely unlikely to be due to chance (thanks to a very large number of comparisons), the detected correlation is still very weak (r=0.15). Moreover, the correlation plots (Supplementary figures S9 and S10 in Wang et al. [2017]) clearly show that this weak correlation is due to geographically distant pairs that are genetically dissimilar (the top right quarter of the plots), and not due to geographically close pairs that are genetically similar. This implies that geographically close pairs do not display correlation between genetic and geographic distances, which means that gene flow between wild and domesticated rice is not detected by this test.

To conclude, genome-wide diversity scans have repeatedly revealed unique diversity patterns in *indica*, *japonica* and *aus* rice, indicating generally different demographic histories. Each of these cultivated rice groups has a set of putative selective sweep regions, with relatively small overlap. The outstanding question of Asian rice domestication concerns the genealogical history of the alleles in this overlap. These alleles are uniform in all cultivated rice and are functionally related to domestication. Currently, there are two competing hypotheses regarding their genealogy: (i) the alleles were selected and fixed in one cultivated group (*japonica* being the usual choice) and subsequently transferred to other (pre)domesticated groups by introgressive hybridization (e.g. Huang *et al*. 2012; Choi *et al*. 2017; Choi and Purugganan 2018); (ii) the alleles existed in different wild populations prior to domestication and were selected multiple times from standing variation in independent domestication processes (e.g. Civáň *et al*. 2015; Civáň and Brown 2017). Since both scenarios are expected to leave similar signatures in the cultivated genomes, it is inherently difficult to decipher the correct answer. Nonetheless, we emphasize that there is currently no conclusive evidence in favour of the introgression hypothesis.

